# Toeholder: a Software for Automated Design and *In Silico* Validation of Toehold Riboswitches

**DOI:** 10.1101/2021.11.09.467922

**Authors:** Angel F. Cisneros, François D. Rouleau, Carla Bautista, Pascale Lemieux, Nathan Dumont-Leblond, Team iGEM ULaval 2019

## Abstract

Synthetic biology aims to engineer biological circuits, which often involve gene expression. A particularly promising group of regulatory elements are riboswitches because of their versatility with respect to their targets, but early synthetic designs were not as attractive because of a reduced dynamic range with respect to protein regulators. Only recently, the creation of toehold switches helped overcome this obstacle by also providing an unprecedented degree of orthogonality. However, a lack of automated design and optimization tools prevents the widespread and effective use of toehold switches in high throughput experiments. To address this, we developed Toeholder, a comprehensive open-source software for toehold design and *in silico* comparison. Toeholder takes into consideration sequence constraints from experimentally tested switches, as well as data derived from molecular dynamics simulations of a toehold switch. We describe the software and its *in silico* validation results, as well as its potential applications and impacts on the management and design of toehold switches.

## 1. Introduction

### 1.1 Riboswitches

All biological systems, be they naturally occurring or synthetic, rely on finely tuned interactions of their components. The precise regulation of these interactions is often critical to proper system functions, and there exist, in nature, many such regulatory mechanisms. A particularly interesting group of regulatory elements are riboswitches - RNA molecules, which typically predominate within the 5’-untranslated region (UTR) of prokaryotic protein coding transcripts and that fold into specific secondary and tertiary structures capable of regulating transcription and translation, thereby optimizing the use of resources (Findeiß et al. 2017). Riboswitches have been observed in bacteria (Winkler, Nahvi, and Breaker 2002), archaea (Gupta and Swati 2019), and in some fungi and plants (Sudarsan, Barrick, and Breaker 2003). They respond to a wide range of stimuli, for instance metabolite concentrations, and their prevalence and versatility in nature makes them attractive for the design of synthetic biological circuits (Mandal and Breaker 2004; Garst, Edwards, and Batey 2011).

Efforts to leverage the potential of riboswitches for synthetic biology have led to several different designs. Out of these, toehold switches have recently been put in the spotlight as a versatile tool with an unprecedented dynamic range and orthogonality (orthogonality meaning that the system is self-contained and has as little spurious effects as possible on other cellular functions) (Green et al. 2014). Toehold switches are single-stranded RNA molecules containing the necessary elements for the translation of a reporter protein: its coding sequence, a ribosome binding site, and a start codon. They fold into a specific hairpin-like secondary structure that blocks the ribosome’s access to its binding site and the first start codon on the RNA strand, therefore preventing translation of the coded protein further downstream (OFF state). The hairpin is designed such that when the toehold riboswitch is in the presence of its DNA or RNA “trigger” sequence, the hairpin unfolds (ON state), hence giving access to the ribosome binding site and the start codon to enable translation (Green et al. 2014) (Figure 1). As a result, the reporter protein can be used to confirm the presence of the trigger sequence in a sample, which opens a wide variety of potential applications for biosensors.

**Figure 1:**
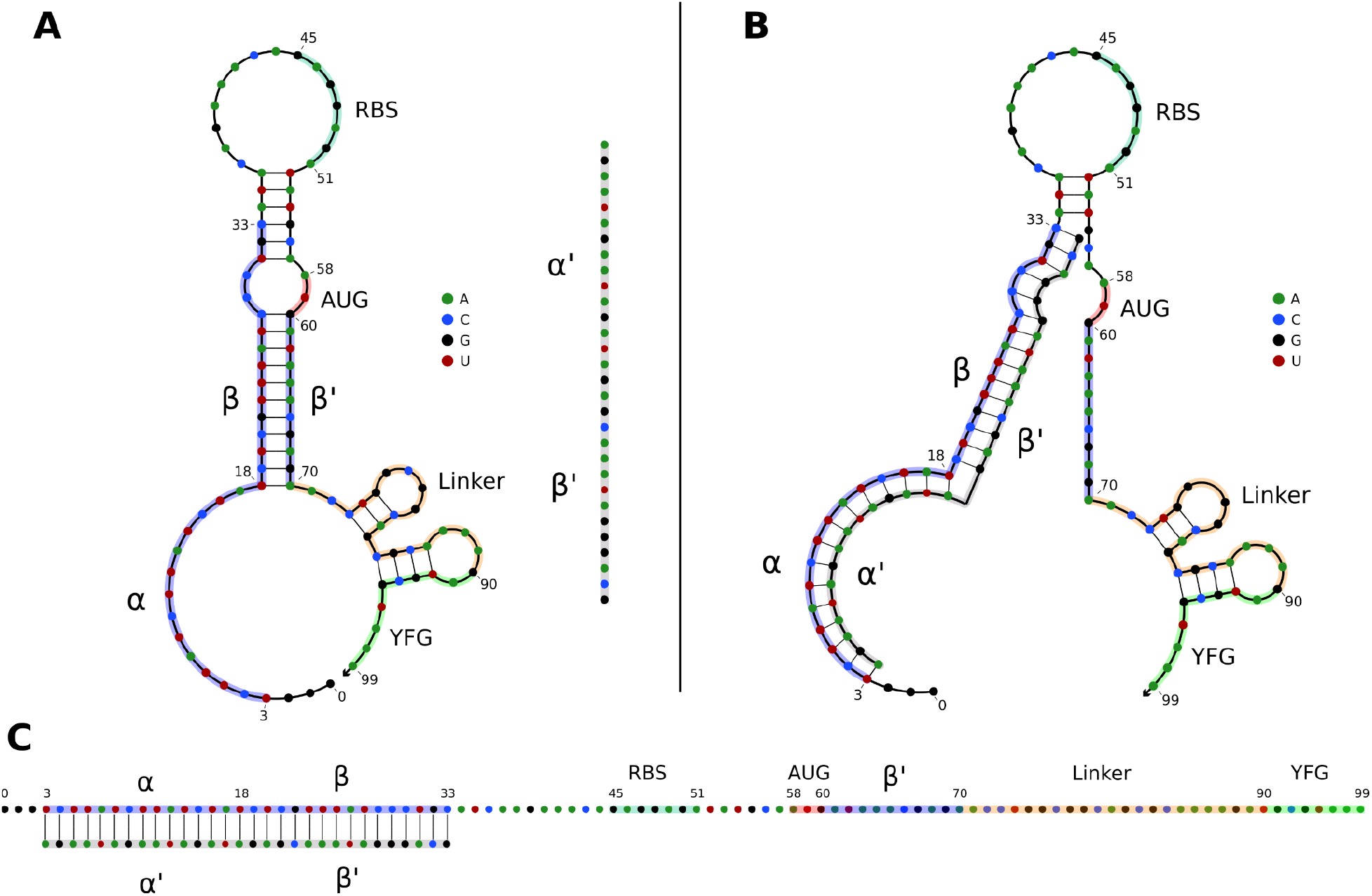
**A)** OFF state of a typical toehold switch. Nucleotides (nt) 3 to 33 (α, β) are complementary to the trigger sequence (α’, β’), nt 45 to 51 are the RBS, nt 58 to 60 are the upstream start codon, nt 70 to 90 are the linker sequence, nt 90 and downstream are part of the regulated gene of interest. The trigger sequence (α’, β’) is shown in grey for reference next to the toehold switch. **B)** Intermediate state of a toehold switch when it first binds to its trigger sequence. **C)** ON state of typical toehold switch, where it is stably bound to its trigger sequence, and translation can occur.

### 1.2 Applications

Despite being a fairly recent technology, toehold switches have already been applied to various fields. Applications include orthogonal systems to regulate gene expression *in vivo* (Green et al. 2014), diagnostic tools for RNA virus detection (ebola (Magro et al. 2017), coronavirus (Park and Lee 2021), norovirus (Ma et al. 2018)), organ allograft rejection detection (Chau and Lee 2020), and even logic gates for gene regulation in synthetic systems (Green et al. 2014, 2017) for pharmaceutical and medical purposes, for example as targets for novel antibiotics (Blount and Breaker 2006) or in gene therapy (Nshogozabahizi et al. 2019). Toehold switch-based technology is highly modulable and cost-effective, making it a very interesting tool to address present and future challenges, and holds great promise in being extendable to numerous and varied purposes.

### 1.3 Design

When the toehold switch is properly designed, the hairpin will natively fold on itself as the RNA is transcribed, following Watson-Crick canonical hydrogen bonds-based pairing. In absence of the trigger sequence, it will be most stable when in its OFF (hairpin/unbound) conformation, therefore preventing spurious activation and translation of the downstream open reading frame (ORF). In presence of the trigger sequence, the higher Watson-Crick homology between the switch/trigger structure than within the switch itself will favor the unfolding of the hairpin (the ON state), allowing for downstream translation.

However, the design of toehold switches is not always straightforward. As proper repression of the downstream ORF relies on the secondary structure to avoid leakage and spurious translation, the sequence of the hairpin structure, and therefore the sequence of the trigger, is critical. Depending on the trigger sequence, many of the regulatory parts of the toehold switch, including the RBS and first start codon, and to a lesser extent, the linker sequence, can interfere with proper folding of the hairpin (Findeiß et al. 2017). There are therefore important sequence constraints to observe when designing good quality toehold switches, in which signal leakage (OFF activity) is minimized, while maximizing protein expression (ON activity) when bound to its trigger. Therefore, studying the molecular dynamics of toehold riboswitches could help identify ways to improve their design.

Over the past few years, leaps and bounds have been made in the field of toehold switch design. Vast improvements have been made on their ON/OFF ratios/fold increase, dynamic expression levels, and signal leakage, and some sites on the trigger sequence have been identified as being key to hairpin folding, but a standardised “best-practice” when designing toeholds is still lacking. Since few high-throughput datasets on experimentally tested toeholds are available, understanding what makes some better than others remains difficult (Green et al. 2014). As of right now, the main limiting factor in the broader applications of toehold technology is the exploratory aspect of designing toehold switches, as well as intrinsic limitations imposed by essential switch elements (Ausländer and Fussenegger 2014).

In 2019, our iGEM team designed a project around the real-life applications of toehold switches. Thus, we looked for available tools that could aid the design of these riboswitches. To the best of our knowledge, the only available tools for the design of toehold riboswitches were the NUPACK design suite (Zadeh et al. 2011) and a tool designed by Team iGEM CUHK 2017 (To et al. 2018). However, these tools have a high entry level difficulty, especially when setting up a methodology and when analyzing the results. To address this, our 2019 iGEM team decided to design an open-source software to make working with toehold switches more accessible, and hopefully allow for broader applications of toehold-based technologies. We created Toeholder, a comprehensive software for toehold design and *in silico* comparison. Toeholder takes into consideration sequence constraints described by Green et al (2014), as well as data derived from our molecular dynamics simulations of a toehold switch. In the present work, we describe the software and its *in silico* validation results, as well as its potential applications and impact on the management and design of toeholds.

## 2. Materials and methods

### 2.1 Molecular dynamics simulations of a toehold switch

Molecular dynamics simulations were performed on a toehold switch from Green et al. (2014) to study the dynamics of its predicted secondary and tertiary structure. We hypothesised that fluctuations in the formation of hydrogen bonds in the hairpin of the toehold switch could lead to spontaneous unwinding of the hairpin, causing the residual OFF signal observed in experiments. As such, we reasoned that studying the dynamics of the structure might provide a broader understanding of the stability of the base pairing in toehold switches.

Sequences from previously designed toehold switches were downloaded from Green et al. (2014). Toehold switch number 1 from table S3 was selected for further modeling because it provided the highest ON/OFF ratio. Its sequence was used to generate a secondary structure with NUPACK (Zadeh et al. 2011) with the rna1995 parameters (Serra and Turner 1995; Zuker 2003; Dirks and Pierce 2003) and a temperature of 37°C. Later, the sequence and the predicted secondary structure were submitted to the RNAComposer online server (Popenda et al. 2012; Purzycka et al. 2015) to obtain a 3D model. The quality of the 3D model was validated with MOLProbity (V. B. Chen et al. 2010) (Table S1). The 3D structure of the toehold switch was introduced in a square water box (146 Å x 146 Å x 146 Å) using the online CHARMM-GUI server (Jo et al. 2008; Lee et al. 2016) with a salt concentration of 0.15 M NaCl. Energy minimization was performed using an NPT equilibration at a constant temperature of 298.15 K. Molecular dynamics simulations were run with the NAMD simulation engine (Phillips et al. 2005) with explicit solvent and periodic boundary conditions for a total length of 40 ns using the CHARMM36 force field and the TIP3P water model.

Molecular dynamics simulations (Supplementary video 1) were analyzed using VMD (Humphrey, Dalke, and Schulten 1996). The stability of the hairpin of the toehold riboswitch was evaluated by measuring the persistence of hydrogen bonds throughout the simulation. The percentage of frames in the simulation in which a hydrogen bond is detected (occupancy) was measured using VMD with a distance cut-off of 3 Å and an angle cut-off of 20°. Hydrogen bonds were classified as either canonical (if they appear in the desired secondary structure) or non-canonical (if they do not).

### 2.2 Designing toehold switches with Toeholder

In parallel to the previous tests, an automated workflow to design and test toehold switches was created to accelerate those processes. The Toeholder software is publicly available on GitHub at https://github.com/igem-ulaval/toeholder. As of publication, it is the first iteration of the program built on the observations of Green et al. (2014). Improvements based on our molecular dynamics simulations remain to be made.

The Toeholder workflow for designing toehold switches is shown in Figure 2. Briefly, Toeholder receives a target gene and other parameters (length of trigger region bound to target, length of trigger in hairpin, reporter gene sequence) as input that will be used to perform a sliding window scan of the target sequence. The sliding window is used to determine the trigger sequence, that is, the complement of the intended target sequence. Afterwards, the sequence that will close the hairpin is added as the complement of the second part of the trigger sequence. The loop and linker regions are taken from the sequence of toehold 1 from table S3 from Green et al. (2014). Once the candidate toehold for that window has been produced, the sliding window advances by one nucleotide. Toeholder produces potential switches for candidates along the entire length of the target gene.

**Figure 2.**
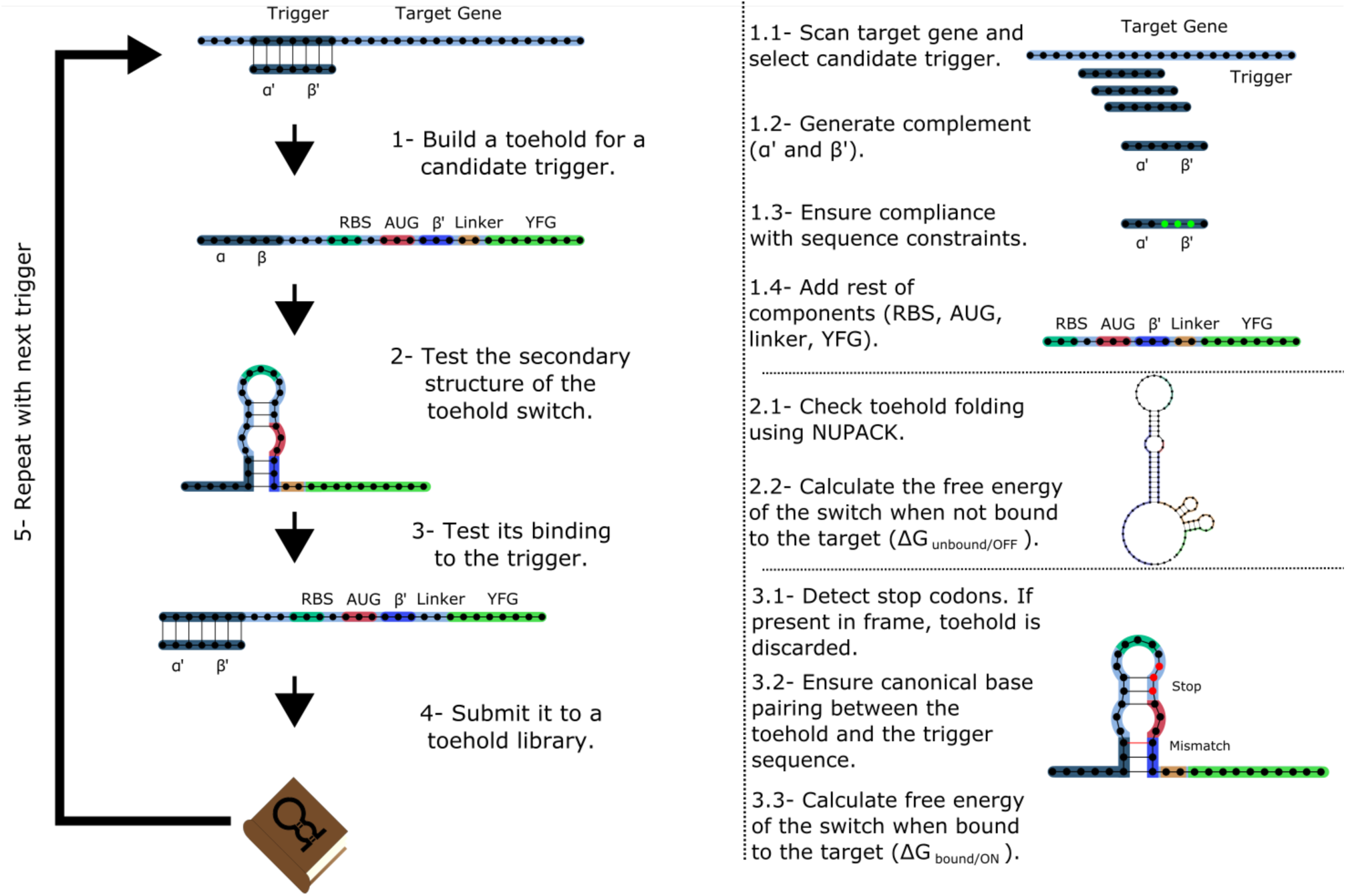
Workflow used by Toeholder to design toehold riboswitches. From a target gene, a sliding window is used to determine candidate triggers and its complementary sequence is used to produce the hairpin. The rest of the elements of the toehold riboswitch are then added to the sequence. The secondary structure, binding energy, and binding accuracy of the toehold riboswitch are then tested *in silico*. Toeholder saves the results and moves the sliding window by one nucleotide to work with the following candidate trigger.

Toehold switches produced by Toeholder are then tested automatically using NUPACK (Zadeh et al. 2011). The minimum free energy secondary structures of the proposed toehold switch and the target mRNA are generated separately, as well as the minimum free energy secondary structure for the proposed toehold switch bound to the target mRNA. The calculated free energies from these three tests are used to determine the changes in free energy (ΔΔG) (Formula 1).

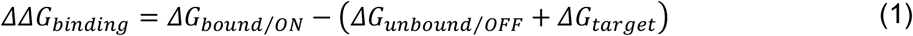

The potential switches with the lowest ΔΔG_binding_ are considered the most likely to offer good performance. Furthermore, the predicted structure of the toehold switch bound to the target mRNA is used to test if the hybridized region is the intended target. Toehold switches that bind perfectly to the intended target are prioritized over those that are predicted to bind partially. The final tests involve looking for stop codons in the region of the toehold switch that would be used for translation, which results in a toehold switch being discarded, as well as ensuring canonical base pairing along the hairpin structure. Finally, only switches which respect suggested forward engineered sequence constraints based on experimental evidence from Green et al. (2014) (2 G:C / 1 A:U base pairing at the bottom of the hairpin, 3 A:U base pairing at the top of the hairpin) are passed to the output.

### 2.3 Validation of Toeholder

Toeholder was created as part of a bigger project, A.D.N. (Air Detector of Nucleic Acids), that was meant to detect pathogenic viruses in the air through a combination of toeholds based biosensors and microfluidics. Therefore, the Toeholder workflow (see section 2.2) was used to design and test *in silico* toehold switches for seven different targets. These targets were selected on the basis of feasibility of our iGEM team working with them in a laboratory (*oxyR* from *Escherichia coli*, two CDS from the Phi6 bacteriophage, an ORF from the bacteriophage PR772) or viruses that can represent health concerns (norovirus, measles virus H1, human alphaherpesvirus 3). The *in silico* characterization of the switches and their production process gave us a substantial validation of the initial workflow. The resulting switches, as well as the accession numbers of the target sequences are detailed in Table S2. Ultimately, the three switches with the lowest ΔΔG_binding_ and perfect matches to their respective triggers for each target were selected and submitted as parts to the iGEM registry. Selecting three candidates per target allows for a greater probability of identifying a successful switch, since our iGEM team was unable to validate them experimentally.

Toeholds were also aligned to several reference genomes to test their predicted specificity and versatility using blastn for short sequences (Camacho et al. 2009). These reference genomes were selected based on the possibility of being present in the same samples as the target in a real application (*Escherichia coli, Homo sapiens*, MS2 phage, PM2 phage, Norovirus, Herpesvirus) and to determine if the trigger sequence of a toehold switch was present in several different measles virus strains (B3, C2, D4, D8, G2, H1).

## 3. Results

### 3.1 Analysis of molecular dynamics simulations

The modeled structure of the toehold riboswitch from Green et al. (Green et al. 2014) remained stable throughout the molecular dynamics simulation (supp. video 1). In particular, the hairpin of the toehold riboswitch did not unwind, which would have led to the unwanted expression of the reporter gene. The most flexible regions of the structure were the two ends of the molecule, as expected, because base pairing in these regions is very limited.

Since the hairpin relies primarily on hydrogen bonds resulting from base pairing, we did a quantitative analysis on hydrogen bonds throughout the molecular dynamics simulation. We found that the number of hydrogen bonds remains relatively stable throughout the simulation (Figure 3A), which is consistent with our observation of the hairpin not unwinding. We then set out to identify the positions in the hairpin that were responsible for the fluctuations observed in the number of hydrogen bonds. We measured the occupancy, i.e. the percentage of frames of the simulation in which the hydrogen bond is observed, of each intended hydrogen bond in the hairpin (Figure 3B). Since base pairing includes multiple hydrogen bonds (two for each A:U pair and three for each G:C pair), each position is represented by the mean of the occupancies of its hydrogen bonds. By comparing the occupancies at each position, we identified the five most stable (hydrogen bonds between nucleotides 19, 21, 22, 32, and 33 and their complements) and the five least stable hydrogen bonds (nucleotides at positions 23, 24, 26, 31, and 36 with their complements) of the hairpin of the simulated toehold switch (Figure 3C). Thus, we hypothesized that GC content at these positions of interest could facilitate hairpin unwinding and contribute to the high ON/OFF ratio of toehold switch 1.

**Figure 3.**
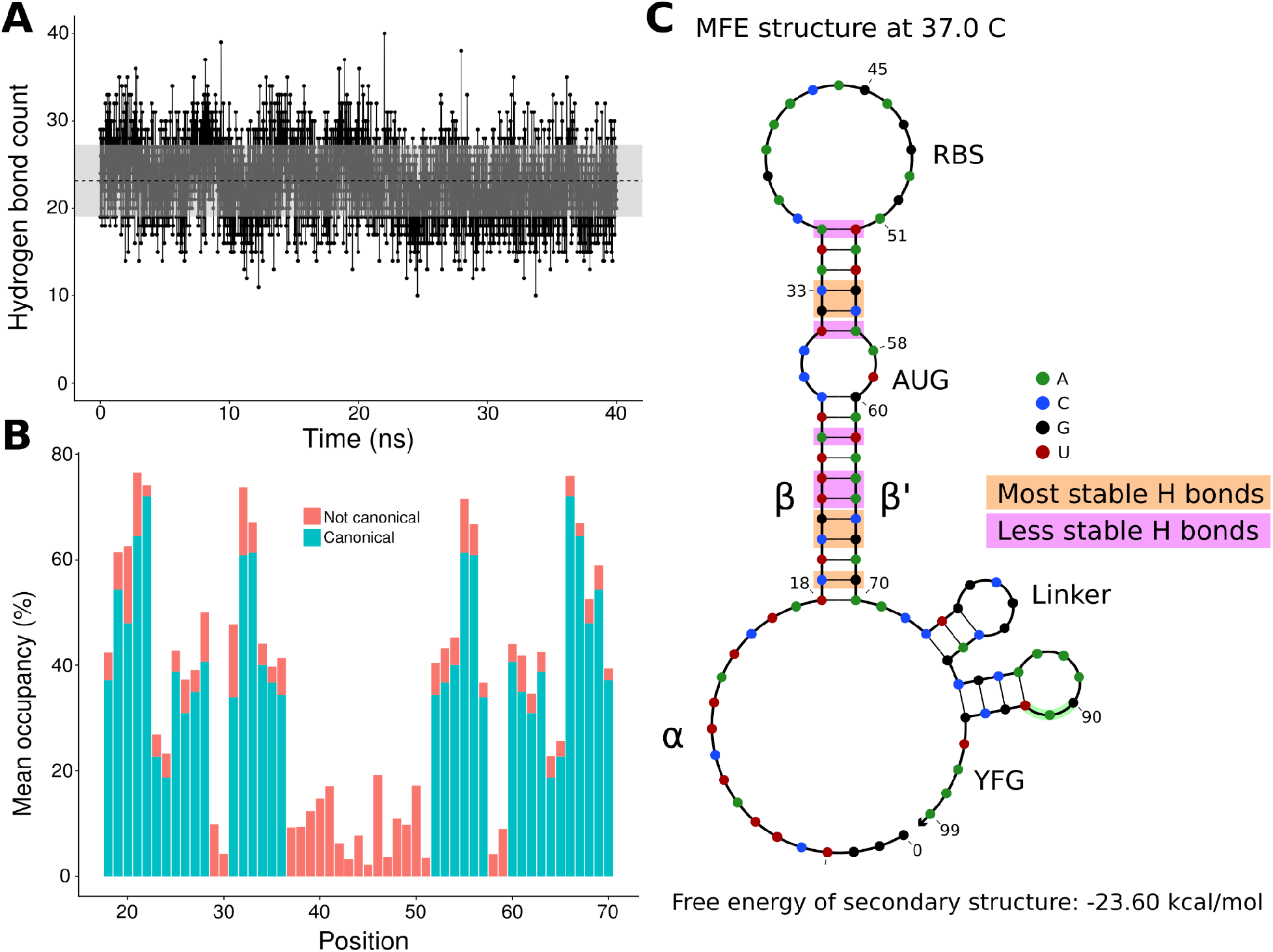
Analysis of hydrogen bonds throughout the molecular dynamics simulation. **A)** Number of hydrogen bonds observed at every time point of the simulation. The black dashed line indicates the mean number of hydrogen bonds, and the shaded region indicates one standard deviation above and under the mean. **B)** Average occupancy of canonical (as determined by the predicted secondary structure) and not canonical hydrogen bonds throughout the molecular dynamics simulation at each position. **C)** Secondary structure diagram showing the positions with the most and least stable hydrogen bonds in the hairpin.

To test the contribution of GC content at these positions of interest to ON/OFF ratio, we reanalyzed the available dataset of 168 first-generation toehold switches from Green et al. (2014). We labeled each of the toehold switches based on the number of positions of interest from the molecular dynamics simulation containing GC, except for position 36 since design constraints require A:U pairing at that position. However, our statistical test (ANOVA with Tukey’s test for honest significant differences) showed that any differences in ON/OFF ratio for toehold switches with GC at the most stable positions (Fig. 4A) or at the least stable positions (Fig. 4B) were not statistically significant. To complement the analysis, we analyzed the distribution ON/OFF ratio based on the combination of GC content at both the most stable and least stable positions but observed that the available dataset underrepresents most of the possible combinations, with no switches sharing the pattern observed in toehold switch 1 of GC at all of the most stable positions and AU at all of the least stable positions (Fig. 4C). Thus, our results suggest that neither the most stable nor the least stable positions could explain the ON/OFF ratio on their own, but we cannot fully confirm the relevance of these positions based on currently available experimental data.

**Figure 4.**
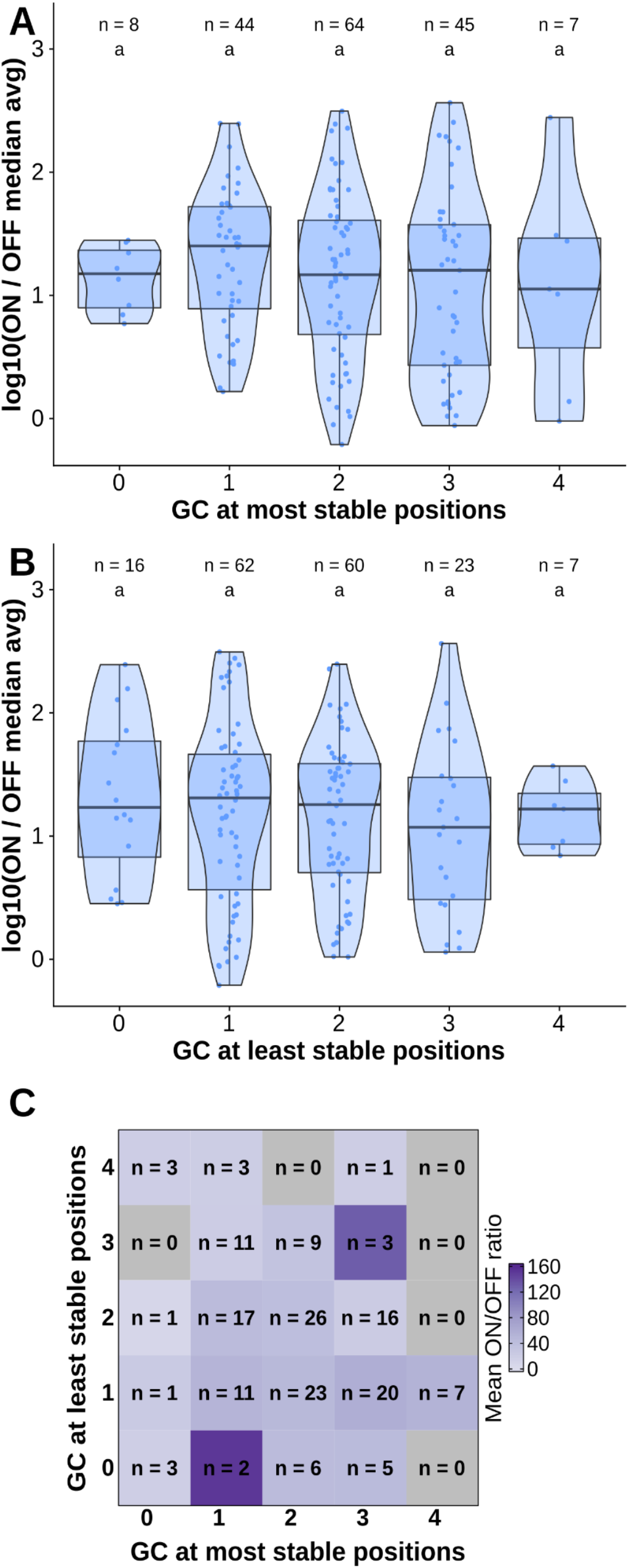
Contributions of GC content at positions of interest from the molecular dynamics simulation. Data from first-generation toehold riboswitches from Green et al. 2014 were used. **A)** ON/OFF ratio for toehold riboswitches based on GC at the most stable positions for the molecular dynamics simulation of the best forward engineered toehold from Green et al. 2014. **B)** ON/OFF ratio based on GC at the least stable positions from the molecular dynamics simulation. **C)** Combinations of GC at the most stable and least stable positions and the mean ON/OFF ratio for each combination. Numbers of toehold riboswitches in each group are indicated.

### 3.2 Validating toehold riboswitches designed by Toeholder

All toehold riboswitches designed by Toeholder were tested *in silico* to evaluate their quality. Here, we show how riboswitches designed with Toeholder for seven different targets scored in our tests.

The first test validates the secondary structure of the riboswitch using NUPACK (Zadeh et al. 2011). Our riboswitches tended to have a similar secondary structure to the one with the highest ON/OFF ratio designed by Zadeh et al. (2011). The average secondary structures for riboswitches generated for each of the seven different targets and the riboswitch from Zadeh et al. (2011) as the reference are shown in table S2. Average secondary structures were generated by taking the most frequent state for each position in the set of sequences for the same target. Importantly, the main hairpin and the smaller one closer to the reporter gene are preserved in these average secondary structures, indicating that toehold riboswitches designed by Toeholder fold into a desirable secondary structure.

The following tests evaluate the predicted binding of the toehold riboswitches to the target. The distributions of ΔΔG_binding_ values for every toehold riboswitch candidate produced for the seven targets are shown in Figure 5A. Since all the ΔΔG_binding_ are negative, the bound state is more stable for all of our riboswitches than the unbound state.

**Figure 5.**
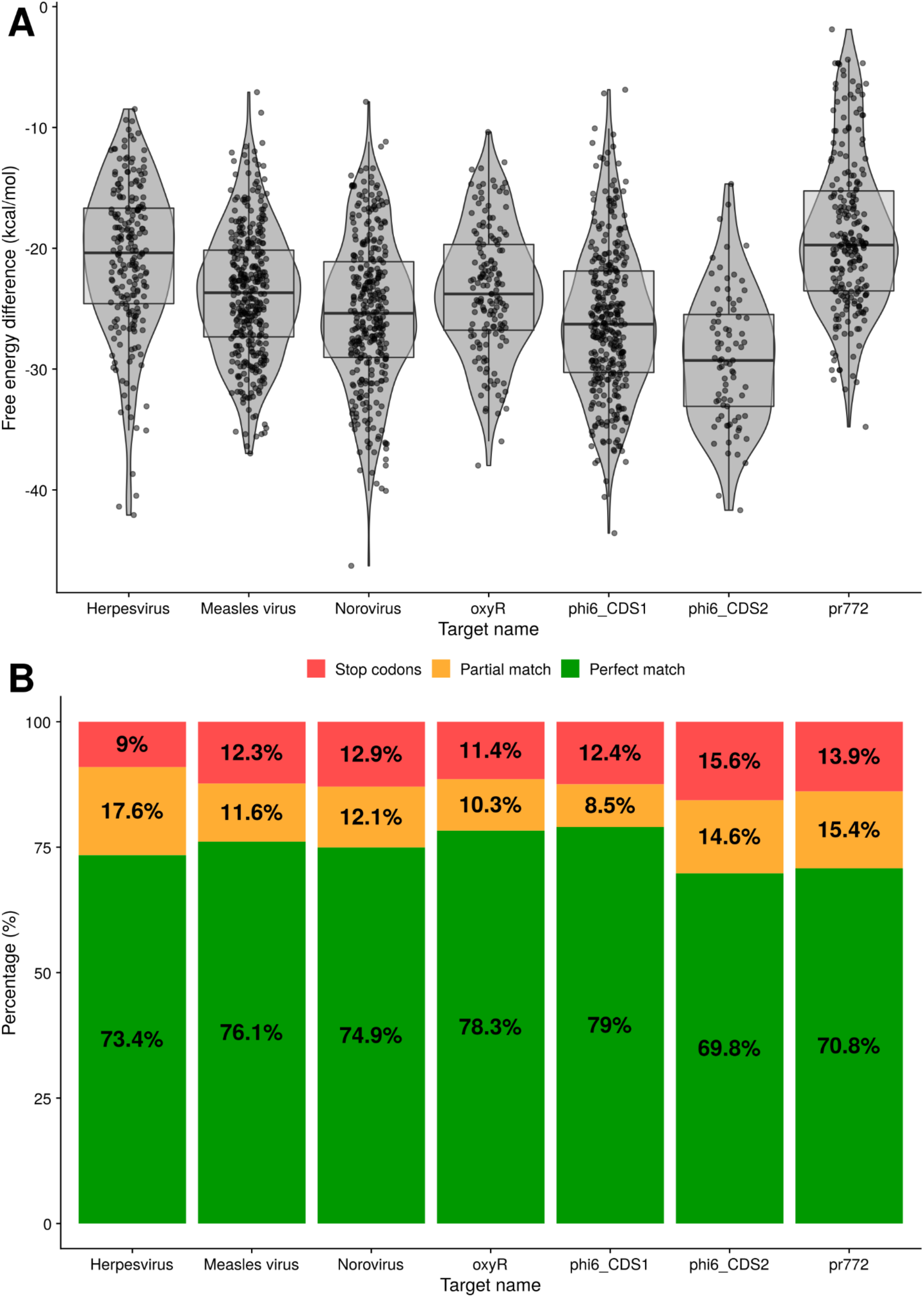
Analysis of binding for toehold riboswitches designed by Toeholder. A) Distribution of free energy differences between the unbound state and the bound state among the number of toehold candidates. B) Classification of toehold riboswitches according to the accuracy with which they bind to their target (imperfect and perfect match) and if they have a stop codon.

Similarly, using the prediction for the bound secondary structure, we can evaluate if each designed toehold riboswitch is predicted to bind to its intended target. Toehold riboswitches were classified as perfect matches if all their positions were predicted to bind to the target and imperfect matches if there was at least one mismatch. As shown in Figure 5B, around 70% of the riboswitches designed for each of our targets are predicted to bind perfectly, even when discarding all the ones that have undesirable stop codons. Thus, our riboswitches would be expected to be able to recognize their targets efficiently.

## 4. Discussion

### 4.1 Toehold switch characterization through molecular dynamics

Molecular dynamics simulations were first performed to get insights into the molecular interactions in the toehold structure. Our results allowed us to identify regions more likely to play an important role in the ability of switches to retain their appropriate secondary structure in the absence of the trigger. The results obtained were in line with the structural description given by Green et al. (2014). The 3D structure of the switch was stable under the conditions it was tested in (0.15M NaCl, 298.15K).

The stability of the hydrogen bonds responsible for this structure were also studied to identify weakpoints that may be worth considering when designing toehold switches. The base pairing of nucleotides at positions 23, 24, 26, 31, and 36 with their complementary sequences fluctuates the most often during the simulation, yet it is critical in preserving appropriate folding and reducing OFF signal. To reduce spurious expression of the reporting protein in absence of the target, it may be useful to favor guanine or cytosine bases in those positions to increase structural stability. Since this may also come at the cost of reduced sensitivity, additional data and *in vitro* tests are required to confirm these assumptions empirically. It is also important to remember that these weaker sites could change for toehold switches with different specifications, such as longer or shorter hairpins. Therefore, further analyses with longer simulations of more switches could help identify the positions of interest for different designs. It should also be noted that the mean occupancies presented in figure 3 were computed on a different number of hydrogen bonds depending on the type of nucleotide (A:U = 2 bonds, G:C = 3 bonds) and that it does not allow for individual characterization of those bonds. However, since only entire nucleotides can be substituted, and not individual bonds, we believe this representation remains useful to identify and consolidate structural weaknesses.

### 4.2 Toeholder conception

In parallel to these experiments, we created Toeholder, an automated workflow for toehold switches design based on sequence requirements defined by Green et al. (2014). The open-source program, that can be run locally or at our web server (https://toeholder.ibis.ulaval.ca/), allows the users to input target sequences and receive a list of potential toehold sequences that have been curated and ranked. As a result, we believe Toeholder will contribute to a reduction of the high entry level difficulty usually associated with this molecular regulator technology.

The output of Toeholder is fully described in the Github repository. Briefly, results are organized in a folder containing copies of the input files, tables summarizing the results for all the toehold switches, and individual subfolders for each of the switches designed. Users would be encouraged to select toehold riboswitches to test experimentally based on the data available (free energy change of binding to the target, whether the toehold is predicted to bind perfectly to the trigger sequence, the desired specificity or versatility depending on matches found in genomes of interest, and the percentage of GC in weaker regions of the hairpin). Once selected, the user can find the full sequence of the riboswitch in its respective subfolder based on its index.

Toeholder also allows users to submit genomes of interest to search for hits of the trigger sequence. This function can be used to evaluate if a riboswitch satisfies the needed requirements of target specificity or universality. For example, we tested for hits of our trigger sequences in the human genome. This allowed us to confirm that the sequences targeted by our toehold riboswitches were not present in the human genome, thus minimizing the possibility of having spurious expression due to the riboswitches interacting with human sequences. On the other hand, we looked for hits in several measles virus strains in order to make sure the trigger sequences were conserved, so that the designed riboswitches would be able to recognize many of the different strains.

The potential improvement in sequence composition found using molecular dynamics have not been added to the program. Yet, due to its open-source nature, these modifications can be easily introduced retroactively, through the Github repository, when more robust data supports the importance of these positions in detection effectiveness. Due to temporal and monetary limitations, we were unable to experimentally assess the importance of these sites. However, since they follow experimentally validated constraints from Green et al. (2014), we believe that the toehold switches produced by Toeholder should operate in a dynamic range similar to that of the forward-engineered switches from this experimental dataset.

### 4.3 Toeholder validation

Toeholder was used as part of our 2019 iGEM project to design switches that could detect phages and bacterial components used for *in vitro* and proof of concept tests, as well as switches for human viruses. Additional tests were run on the outputs of the designs to validate the program. First, the secondary structure of all the riboswitches candidates for the seven targets were computed using NUPACK and all of them presented a similar structure to the one we characterized from Green et al. (2014). Therefore, we expect them to behave in a similar way *in vitro*. Their free energies were also recomputed and are presented in figure 5A. All switches have a negative energy that predicts they should favour the bound state to the target. In addition, of all the candidate switches produced, around 70% and up were a perfect match to the target, meaning Toeholder effectively suggested switches that would theoretically recognize their appropriate target. Altogether, the software consistently produced candidate switches that are within the defined sequence and structural restrictions and that should recognize their target, all of it in an easy-to-use format.

### 4.4 Comparison with different approaches

Although the study of riboswitches is currently somewhat limited to proof-of-concept studies, *in silico* approaches have been widely explored for prediction of riboswitches performance both from sequence information alone (Barrick 2009; Nawrocki, Kolbe, and Eddy 2009) and structural features (Barash and Gabdank 2010). However, despite the many possibilities and applications that Toehold switches offer, far fewer studies have focused on the *in silico* design of these tools specifically ((Zadeh et al. 2011), (To et al. 2018)). The lack of high-throughput datasets on experimentally tested toeholds makes it difficult to understand what affects their performance and how it can be improved. Therefore, our open-source software, in addition to allowing the high-throughput effective design of Toehold switches, provides a global idea of their dynamics and operation. Besides its simplicity in terms of design, we have provided an *in silico* validation, which ensures an effective and working design.

### 4.5 Limitations

The limitations of Toeholder reside in its fully *in silico* approach. Our computations may overlook sequence requirements that could only be discovered by extensive *in vitro* experiments. Very few data sets of such nature are currently available, and we were unable to complete these experiments on the switches we designed for the 2019 iGEM competition, due to time constraints. Questions also remain on the optimal physicochemical conditions to use toehold switches. Our *in silico* models and validation use standard conditions, in part limited by the programs, that may not reflect the way switches may want to be used. Certainly, the conditions are critical in the control of these tools since natural riboswitches can detect concentrations of small ligands (reviewed in (Findeiß et al. 2017)), but are sensitive to changes in temperature (Narberhaus 2010) or pH-value (Nechooshtan et al. 2009) which can be a limitation if conditions are no longer controlled, reducing their potential applications in very different systems or in extreme conditions. However, toehold switches address some other limitations of earlier riboregulator designs as low dynamic range, orthogonality, and programmability, since these RNA-based molecules exhibit more kinetically and thermodynamically favorable states by incorporating linear - linear interactions instead of loop-loop and loop-linear interactions (Green et al. 2014). This reflects the need for high throughput experimental screening to accompany *in silico* studies such as this one. However, our software provides a first step to facilitate high-throughput toehold switch design, production, and testing. Future studies could use it as a steppingstone to provide more in-depth characterization of these promising molecular regulators and therefore, to overcome their limitations.

### 4.5 Applications and 2019 iGEM project

Due to their adaptability, toehold switches offer great possibilities of applications. As part of the 2019 iGEM competition, we presented the project A.D.N. (Air Detector for Nucleic acids), which takes advantage of this technology to create a biosensor that detects airborne pathogens (see Team iGEM ULaval 2019 wiki: https://2019.igem.org/Team:ULaval). Riboswitches were designed as the sensing component of a modular device designed to sample air, extract ribonucleotides, and prepare samples via microfluidics, as well as perform detection through fluorescence measurements. The combination of toehold switches with optical detection offers great practicality and target versatility.

## 5. Conclusions

The development of synthetic biology and the numerous molecular systems requires the parallel coupling of bioinformatics tools that facilitate their easy handling and implementation. Our open-source software, Toeholder, aims to facilitate the automated *in silico* design of toehold riboswitches and the selection of switch candidates for a target gene. Furthermore, by using molecular dynamics simulations, we identified the nucleotides in the hairpin of a reference toehold switch whose hydrogen bonds fluctuate the most. These could be potential targets to modify when polishing the design of these riboswitches. Increasing switches efficacy will likely contribute to their integration into broader applications of toehold-based technologies.

## 6. Acknowledgments

Special thanks to all the other members of the iGEM ULaval 2019 team (Catherine Marois, Elodie Gillard, Florian Echelard, Guillaume Fournier, Jean-Michel Proulx, Julien Roy, Karine Bouchard, Lucas Germain, Marianne Côté, Martine Voisine, Nastaran Khodaparastasgarabad, Ahmed Mataich), including our professors and mentors (Hélène Deveau, Michel Guertin, Steve Charette). Team iGEM ULaval 2019 would also like to thank our partners who funded this project and covered participation fees for the competition (https://2019.igem.org/Team:ULaval). We would also like to thank Peng Yin and Alexander Green for kindly sharing additional raw data from their experimentally tested library of toehold switches. Funding sources had no direct involvement in study design or publication of this project.

## 7. Data availability

All data are available in the Supplementary materials.

DOIs:

Toeholder tool: https://doi.org/10.5281/zenodo.7304556

Toeholder data and Scripts: https://doi.org/10.5281/zenodo.7304525

## 8. Declarations of competing interests

None.

## 10. Supplementary data

Supplementary video 1: https://doi.org/10.5281/zenodo.7418392

## Notes

https://github.com/igem-ulaval/toeholder

### Competing Interest Statement

The authors have declared no competing interest.

### Summary of Updates

Adjusted text, added new figures

